# A two-phase plant-soil feedback experiment to explore biotic influences on *Phragmites australis* invasion in North American wetlands

**DOI:** 10.1101/2021.11.12.468454

**Authors:** Sean F.H. Lee, Thomas J. Mozdzer, Samantha K. Chapman, M. Gonzalez Mateu, A. H. Baldwin, J. Adam Langley

**Affiliations:** Department of Ecology and Evolutionary Biology, Tulane University, New Orleans, Louisiana 70118, United States; Department of Biology / Center for Biodiversity and Ecosystem Stewardship, Villanova University, Villanova, Pennsylvania 19085, United States; Department of Biology, Bryn Mawr College, Bryn Mawr, Pennsylvania 19010, United States; Department of Crop and Soil Science, Oregon State University, Corvallis, Oregon 97331, United States; Department of Environmental Science, University of Maryland, College Park, Maryland 20742, United States

**Keywords:** plant-soil feedbacks, *Phragmites australis*, soil feedbacks, *Spartina patens*, invasive plants, soil microbial community

## Abstract

Plants can cultivate soil microbial communities that affect subsequent plant growth through a plant-soil feedback (PSF). Strong evidence indicates that PSFs can mediate the invasive success of exotic upland plants, but many of the most invasive plants occur in wetlands. In North America, the rapid spread of European *Phragmites australis* cannot be attributed to innate physiological advantages, thus PSFs may mediate invasion. Here we apply a two-phase fully-factorial plant-soil feedback design in which field-derived soil inocula were conditioned using saltmarsh plants and then were added to sterile soil mesocosms and planted with each plant type. This design allowed us to assess complete soil biota effects on intraspecific PSFs between native and introduced *P. australis* as well as heterospecific feedbacks between *P. australis* and the native wetland grass, *Spartina patens*. Our results demonstrate that native *P. australis* experienced negative conspecific feedbacks while introduced *P. australis* experienced neutral conspecific feedbacks. Interestingly, *S. patens* soil inocula inhibited growth in both lineages of *P*. australis while introduced and native *P. australis* inocula promoted the growth of *S. patens* suggestive of biotic resistance against *P. australis* invasion by *S. patens*. Our findings suggest that PSFs are not directly promoting the invasion of introduced *P. australis* in North America. Furthermore, native plants like *S. patens* seem to exhibit soil microbe mediated biotic resistance to invasion which highlights the importance of disturbance in mediating introduced *P. australis* invasion.

## INTRODUCTION

Plant-soil feedbacks (PSFs) occur when a plant cultivates a unique soil microbial community and affects the growth of the host plant either positively (positive PSF) or negatively (negative PSF) (Bever 1994, Bever et al. 1997). A growing body of evidence suggests that PSFs can influence the success of invasive plants through direct positive PSFs experienced by the host plant or by negatively affecting the growth of competing plants (Sabelis et al. 2001; Colautti et al. 2004; Mangla & Callaway 2008; Beckstead et al. 2010; Yang et al. 2013; Gundale et al. 2014). Wetlands are disproportionately impacted by invasive plants, yet PSF studies involving wetland plants are lacking as nearly all studies exploring PSFs in invasive plants have been conducted in upland ecosystems.

The European lineage of *Phragmites australis* (Cav.) Trin. ex Steud. (hereafter referred to as introduced *P. australis*) is one of the most prevalent invasive wetland plants in North America. It has spread from the northeastern US to span nearly the entire eastern seaboard, reaching west to the Great Salt Lake, over last half century (Saltonstall 2002). Along with altering ecosystem processes in native wetlands, the invasion of introduced *P. australis* has displaced native wetland plants such as *Spartina patens* and has nearly replaced the native conspecific *P. australis* haplotype “F” in the New England (Windham & Lathrop 1999; Windham & Ehrenfeld 2003; Saltonstall 2003; Silliman & Bertness 2004; Meadows & Saltonstall 2007). The success of introduced *P. australis* in North America contrasts its diminishing abundance in its native range of Eurasia. In western and southern Europe, large die-offs have been referred to as “reed die-back syndrome” (van der Putten 1997; Cerri et al. 2019). In China, invasive *Spartina alterniflora* is displacing *P. australi*s in wetlands it once dominated which is the opposite pattern we observe in North America, where introduced *P. australis* is replacing *S. alterniflora* in high marsh areas (Chambers et al. 1999; Minchinton & Bertness 2003; Vasquez et al. 2006; Zhang et al. 2017^1^). The distinct growth patterns of introduced *P. australis* across a wide variety of habitats in North America compared to Eurasia suggest that variables outside of innate physiology, perhaps biotic interactions such as PSFs, may in part explain the extreme variability in performance.

PSFs may facilitate early establishment survival and dominance of introduced *P. australis*. Several recent studies suggest that *P. australis* soil communities aid in seedling establishment of introduced *P. australis* by protecting and promoting growth in early-stage invasions. For example, previous work suggests that several components of introduced *P. australis*-associated soil communities may have growth-promoting or protective effects. Dark septate endophytes associated with introduced *P. australis* have recently been linked to increasing competitive ability in stressful salinities (Gonzalez Mateu et al. 2020). Other fungal endophytes found in introduced *P. australis* rhizomes can confer protection against pathogens that cause early mortality in seedlings as well as promote germination and growth (Kowalski et al. 2015; Shearin et al. 2018; White et al. 2018). Along with evidence of introduced *P. australis* soil communities facilitating growth, these soil communities also may negatively affect growth of competing plants. Pathogenic *Pythium* oomycetes isolated from introduced *P. australis* soil communities were more harmful to native plant seedlings than isolates taken from native *P. australis* (Nelson & Karp 2013; Crocker et al. 2015). Introduced *P. australis* soil communities can stunt the growth of native *Spartina alterniflora*, while the native *P. australis* soil communities promote growth of native *Spartina alterniflora* (Allen et al. 2018). Thus, pathogen spillover from introduced *P. australis* has the potential to promote the spread of introduced *P. australis* into native wetlands.

The knowledge of inter-lineage and interspecific PSFs is lacking in introduced *P*. australis. We are aware of only one PSF study involving *P*. australis, and it explores intraspecific PSFs among various *P. australis* lineages and *P. australis* soil community effects on *S. alterniflora* (Allen et al. 2018). As an increasing body of evidence supports the importance of seedling recruitment in both establishing new stands of introduced *P. australis* as well as expanding already existing stands, we need to know more about the effects of PSFs on inter-lineage, intraspecific, and interspecific interactions in introduced *P. australis*.

In this study, we explore how PSFs influence the growth of introduced *P. australis* and the native wetland plant *S. patens*. By exploring the direction and strength of inter- and intra-specific PSFs in both these plants we can replicate a typical introduced *P. australis* invasion into a northeastern US wetland. In addition, we test inter-lineage PSFs using native and introduced *P. australis* as well as the variation in PSF effects of native versus introduced *P. australis* soil communities on *S. patens*. We perform a fully-factorial 2-step PSF study to reveal unique PSF interactions by testing every possible PSF combination. The evidence that introduced *P. australis* cultivates soil communities that promote its growth (Kowalski et al. 2015; Shearin et al. 2018; White et al. 2018; Gonzalez Mateu et al. 2020) while eliciting negative growth effects in other species (Nelson & Karp 2013; Crocker et al. 2015; Allen et al. 2018) leads us to hypothesize that (H1) soil communities from introduced *P. australis* would elicit positive/neutral conspecific PSFs (i.e. promote introduced *P. australis* growth) but elicit negative heterospecific PSFs in the native plants native *P. australis* and *S. patens* (i.e. decrease growth in native plants). The decline of native wetland plants in the face of introduced *P. australis* invasion led us to hypothesize that (H2) soil communities from native plants (native *P. australis* and *S. patens*) would elicit positive heterospecific PSFs in introduced *P. australis* (i.e. promote introduced *P. australis* growth) but cause negative conspecific PSFs (i.e. decrease growth in conspecific plant). Addressing these hypotheses will clarify how interspecific and intraspecific PSFs interact in an introduced *P. australis* invasion and help us to determine the possibility that PSFs play a role in facilitating introduced *P. australis* spread into northeastern US wetlands.

## METHODS

### 2-step plant-soil feedback design

In this study we performed a 2-step plant-soil feedback design to study the effects of intraspecific/interspecific PSFs on the growth of introduced *P. australis*, native *P. australis*, and the native wetland plant *S. patens*. In the initial conditioning phase, we take field collected vegetative propagules from all three plants and grow them in sterilized media inoculated with field soil associated with the specific plant (i.e. *S. patens* propagule with *S. patens* inocula). Introduced *P. australis* and *S. patens* each had 12 replicated conditioning pots while *S. patens* had 11 replicated conditioning pots (Fig. 1). After the conditioning phase the conditioned soil was used to inoculate the feedback phase in which all 3 plants were grown in each soil the live soil inocula along with sterilized controls in a fully-factorial design. Each conditioning pot inocula was replicated 2x (2 * 12 = 24 introduced *P. australis*, 2 * 12 = 24 native *P. australis*, and 2 * 11 = 22 *S. patens*, 24 + 24 + 20 = 70 live inoculates per plant species) and used to inoculate the 3 plant species (70 live inocula * 3=210 live inocula pots) within each of the plant species along with 13 sterilized pots within each species (13 replicates x 3 species = 39 sterilized pots), thus the total number of feedback pots is n=249 (Fig. 1). Biomass derived in the feedback phase was used to assess PSFs in each of the plant species.

**Fig.1.**
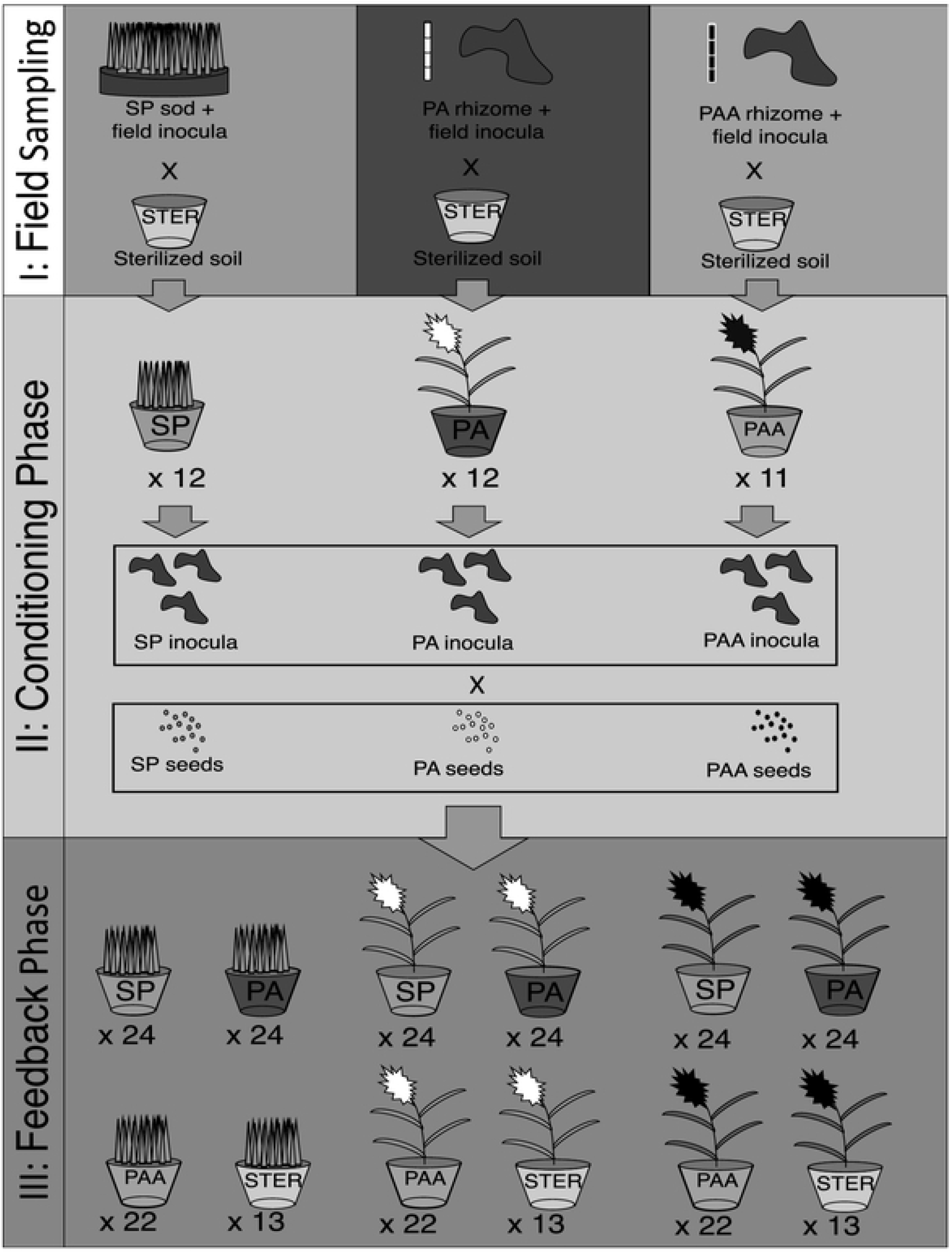
2-step plant-soil feedback design. Each pot represents a different soil inocula x species combination. ^a^SP pots represent *Sparlina patens* inocula, and the spiked shoots and grey seeds representing the *Sparlina patens* plant. ^b^PA pots represent introduced *Phragmiles australis* soil inocula and the white inflorescences/seed represent the introduced *Phragmites australis* plant. ^c^PAA pots represent native *Phragmites australis* soil inocula and the black inflorescencestseed represent the native Phragmiles australis plant. ^d^STER pots represent the stcrilized soil inocula.

### Conditioning phase

#### Field sampling

During the spring of 2017, we collected rhizome fragments from native and introduced *P. australis* stands at King’s Creek (38°46’26.23”N; 75°58’32.38”W), an oligohaline creek that feeds into a section of the Choptank River (0.1 – 3 ppt) in Providence Landing, MD (Table 1). An individual soil/rhizome core (20.3 cm diameter x 30.5 cm depth) were taken 1 m from 6 individual extant introduced *P. australis* shoots, and the same number of cores was collected from an adjacent native *P. australis* stand. A 61 cm x 61 cm x 30 cm (20.3 cm soil depth) sod mat of *S. patens* sods was sourced from the Global Change Research Wetland (GCREW) site at the Smithsonian Environmental Research Center (SERC) in Edgewater, MD (38°53’12.9”N 76°32’24.9”W), a primary tidal creek on the Rhode River (4 - 15ppt) (Table 1).

**Table 1.**
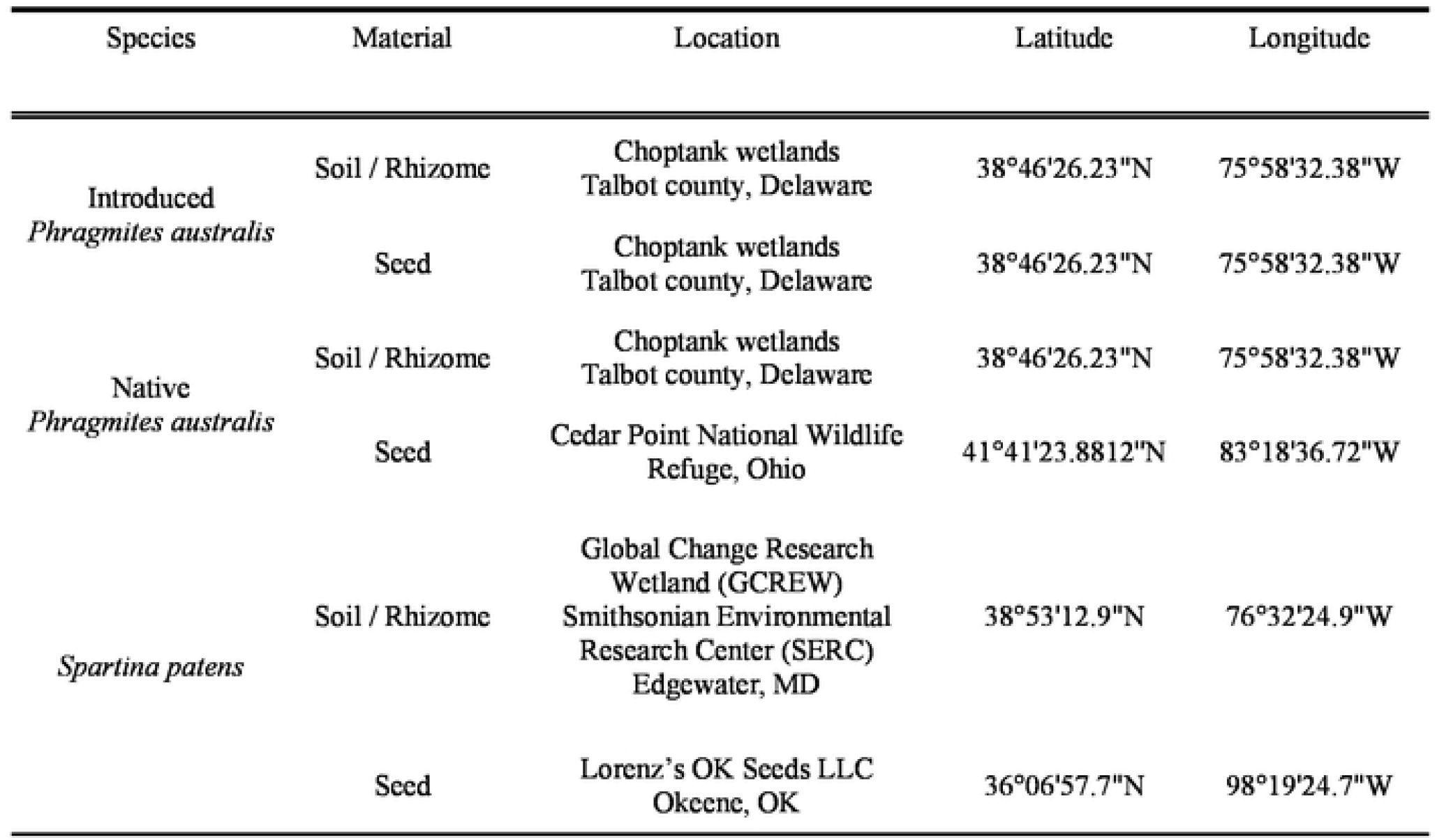
Sampling locations of seed, rhizome, and soil inocula used in the growth experiment

#### Greenhouse design

The “conditioning phase” of this study was performed to ensure that the soil microbial community present in the inocula used to inoculate the subsequent “feedback phase” is representative of the soil community influenced by the target plant species. The six soil/rhizome cores taken from the native and introduced *P. australis* populations were processed to remove rhizome fragments from the soil. The soils from the 6 cores of the introduced P. australis populations and those from the native populations were bulked and used as “field inocula” to inoculate the respective species conditioning pots (i.e. introduced *P. australis* inocula for introduced *P. australis* and native *P. australis* inocula for native *P. australis*). We decided to bulk the soil cores at each site to ensure that the soil community introduced into the conditioning phase was representative of the site, and by sampling from multiple cores within a site we can capture a wider array of plant-associated microbes. The rhizomes of introduced and native *P. australis* were extracted from the soil cores, washed, and divided into 2-3 node segments for planting. The rhizomes were planted 8 cm below the surface of the soil in 2.4 L pots (16.5 cm high x 15.2 cm diameter) filled with a mixture of 900 g of dry sterilized *Sphagnum spp*. moss peat (autoclaved at 121°C for 30 minutes at 30 psi) was mixed into 100 g “field inocula”. Establishment of the *S. patens* conditioning pots followed the same planting procedure as the native and introduced *P. australis* conditioning pots, *S. patens* sods were cut into 5 × 5 × 15 cm sections and used as both soil inocula and propagule source, 900 g of dry sterilized *Sphagnum spp*. moss peat was added to the pot as well. Each conditioning pot contains the focal plant species planted as either rhizome (introduced & native *P. australis*) or as a sod plug (*S. patens*) and the media mix containing conspecific field inocula. A total number of 12 introduced *P. australis*, 11 native *P. australis*, and 12 *S. patens* pots (Fig. 1) were conditioned from April 2017 until December 2017 at the Villanova University Greenhouse (Villanova, PA, USA). Pots were placed in individual reservoirs to avoid microbial cross-contamination and top watered using freshwater every other day until the water reservoir was filled. Although these soil communities were taken from oligohaline sites, we watered with freshwater to avoid confounding variations in salinity. Even so, the soil salinity of the pots at the end of the conditioning phase pots ranged from 2-10 ppt and did not vary consistently among source locations, but instead likely reflected the variability in evapotranspiration among pots. Average daily light intensity (63.8± 8.3 lum m^-2^) and temperature (25.3± 0.1° C) was recorded using the Onset HOBO Pendant^®^ Temperature/Light 8K Data Logger (Onset, Bourne, MA). A subsample of 200-300 g of rhizosphere soil was collected from each of the samples and stored at 4°C (INVAM).

### Feedback phase

#### Seed sourcing

During the fall of 2017, we collected native and introduced *P. australis* inflorescences from the same set of stands at King’s Creek (38°46’26.23”N; 75°58’32.38”W) that we had collected our rhizomes and soil (Table 1). A total of 28 seed-containing introduced *P. australis* inflorescences were collected. Native *P. australis* seed heads were also collected from the stand at King’s Creek but yielded no viable seeds, so 6 seed-containing native *P. australis* inflorescences were sourced from Cedar Point National Wildlife Refuge along Lake Eerie in Northern Ohio (41° 41’ 23.8812’’ N; 83° 18’ 36.72’’ W) in mid-November (Table 1). *S. patens* seeds were sourced commercially (Lorenz’s OK Seeds LLC, Okeene, OK) (Table 1). Native and introduced *P. australis* seeds were extracted from the inflorescences by rubbing the seed heads in-between two gloved hands over a folded sheet of construction paper. Once the seed heads were well masticated the chaff was blown off, leaving only the naked seeds. All seed were pooled and homogenized within each haplotype/species. Logistical issues prevented us from obtaining native *P. australis* and *S. patens* seed from the same populations used in the soil sampling. Native *P. australis* seed heads were collected during the fall 2017 season at the King’s Creek site, however no viable seeds were found on the inflorescences, which suggests we had missed period of seed drop in the native haplotype. The same reason inhibited us from obtaining *S. patens* seed from the GCREW site at SERC and we were forced to purchase seed from a seed supplier.

#### Greenhouse design

Introduced *P. australis*, native *P. australis*, and *S. patens* seeds were surface sterilized using 70% ethanol (3-minute agitation wash) followed by 3% bleach (3-minute agitation wash) after which the seeds were rinsed with ddH_2_O and incubated at room temperature overnight (Kowalski n.d.). The procedure was repeated the next day with the only change being the bleach wash was now 20 minutes. All seeds were then plated on 2% agarose gel and subject to a 16/8 hr diurnal light regime with a 30/15°C day/night temperature to encourage germination. Plates were checked each day to remove molded seeds. The germinated seedlings were then potted into the soil mixtures at 1-cm below the soil surface. The three plant species (introduced *P. australis*, native *P. australis*, and *S. patens*) were grown in soil (700 g of sterilized *Sphagnum spp*. peat moss) inoculated with soil inocula (150 g) from every conditioning-phase pot with two replicates per conditioning phase pot (n=24 introduced *P. australis, n=*22 native *P. australis*, and n=24 *S. patens*) (Fig. 1). There were 70 seedlings of each plant (introduced *P. australis*, native *P. australis*, and *S. patens*) grown in each of the conditioning phase inocula as well as 13 sterilized pots per species (800 g of sterilized *Sphagnum spp*. peat moss and no conditioning phase soils), yielding a total of 248 treatment pots (242 by the end of the trial, Fig. 1). All plants were grown in 1-L pots (6.35 cm diameter x 35.6 cm depth, D60L “Deepot Tree Pots”, Greenhouse Megastore Danville, IL. Each pot was placed in a 946-mL plastic container (11.5 cm diameter x 13.9 cm depth) which served as a water reservoir so that the water table was 21.7 cm below the soil line when filled. All plants received 10 g of Osmocote 15-9-12 slow-release fertilizer in the beginning of March, April, and May (Scotts Co. LLC). All pots were top watered every other day until the water reservoir was filled although overfilling did occur, and pots were not drained throughout the feedback phase. The light and temperature conditions were the same as in the conditioning phase. Because of the short photoperiod during the winter months, an amendment of 3 hours of light from overhead growth lamps was included for the first 8 weeks of seedling growth. Seedling mortality was recorded throughout the length of the experiment and dead seedlings were replaced up until 4/01/2018 (17^th^ week); thereafter pots with dead plants were omitted from the study. Plants were grown from mid-December 2017 and the experiment concluded in late-August 2018.

### Shoot emergence and elongation rate

The rate at which new shoots, shoot emergence (# of new shoots day^-1^), were produced, and rate at which “stand” height grew, shoot elongation rate (mm of growth day^-1^), were measured for each species. Starting in April 2018 total live shoots were counted in each pot and the height of the three tallest shoots was taken. These data were recorded at two additional time points, one in May and one in June. Subsequent measurements were impeded by the increased risk of lodging as pots became denser and taller. Using these data, we calculated the individual slopes of shoot emergence and shoot elongation for each pot.

### Individual plant-soil feedback

To determine the strength and direction of plant-soil feedbacks, total biomass was normalized between within each of the plant species (introduced *P. australis*, native *P. australis*, and *S*. patens) between all soil conditioning treatments (introduced *P. australis*, native *P. australis, S*. patens, and sterilized soil). Nine pairwise plant-soil feedback indices were estimated, 3 for each plant species including two heterospecific plant-soil feedback metrics (PSF_away_) as well as one sterilized plant-soil feedback metric (PSF_ster_) using the methods of Reinhart (2012). For example, we calculated two PSF_away_ values for introduced *P. australis*, one indicating PSF in *S. patens* conditioned soil versus conspecific soil and one indicating PSF in native *P. australis* soil versus conspecific soil. Thus, the target soil conditioning species PSF_away_ varied depended on plant species identity. Soil feedback would be calculated using the formula, PSF = ln(X_C_/X_H_), where X_C_ represents the mean dry biomass of plant species “X” grown in conspecific soil and X_H_ represents the mean dry biomass of plant species “X” grown in a heterospecific or sterilized soil. By observing direction and magnitude of PSFs in each of the plant species we evaluated the soil conditioning effects experienced by each plant species. Positive PSF_away_ would indicate that a plant experiences positive PSFs, growing better in conspecific conditioned soils than in heterospecific conditioned soils. Negative PSF_away_ would indicate that a plant experiences negative PSFs, growing better in heterospecific conditioned soils than in conspecific soils. PSF_ster_ indicates overall soil biota effects and can indicate whether pathogens or mutualists drive PSFs. A plant that displays a negative PSF_ster_ has a conspecific PSF dominated by pathogens while a plant with positive PSF_ster_ has a conspecific PSF dominated by mutualists.

### Substrate-induced respiration (SIR)

We performed substrate-induced respiration assays as an estimate of active microbial biomass and overall activity (Fierer et al. 2003). In July 2018 (∼ 2 months before harvest) soil samples were taken from the feedback growth pots, 8 samples per species x soil conditioning treatment, and 4 samples per species for sterilized soils (n=84). Approximately 10 g of wet soil was taken from the top 4 cm of each pot, making sure to extract soil closely associated to the rhizomes and fine roots. Soil was placed in 55 mL glass headspace vials with rubber septa. Two gas samples were taken in 30-minute intervals for wet soil without the addition of yeast extract broth to establish a baseline respiration rate without substrate. CO_2_ concentration was determined using an infrared gas analyzer (Licor Model LI-7000, LI-COR Biosciences, Lincoln, NE, USA) configured for small sample injection with a six-port valve and a stream of N_2_ as the carrier. Ten ml of yeast extract broth (Sigma-Aldrich, St. Louis, MO, USA) (30 g mL^-1^) was added to each vial containing the wet soil (Barreto et al. 2018). Yeast extract broth was added to provide an excess of labile carbon and nutrients which allowed for quantification of maximal metabolic activity. After the broth was added, headspace was sampled four times roughly at 30-minute intervals to calculate SIR rates. After the final headspace sample, the caps of the septum vials were removed, and the soil slurry was placed in a 50°C oven to dry for 48 hr. Soil mass per vial was calculated by subtracting (vial + dried broth + soil) - (vial + dried broth). Rates were expressed as μg C-CO_2_ g soil^−1^ h^−1^ (Barreto et al. 2018).

### Aboveground & belowground biomass

Feedback phase harvest was commenced over a week starting in late August of 2018. Aboveground biomass was clipped at the soil surface and shoots and leaves were oven dried at 65°C to constant mass to obtain dry biomass measurements. The soil and root clumps remaining were removed from the pots and 10-g soil samples were obtained and stored in 20-mL scintillation tubes from soil clinging to the fine-roots of each of the plants, the soil samples were immediately flash frozen in liquid nitrogen and stored at -80°C. A 5-cm long lateral fine root fragment was taken from each root clump and preserved in 70% ethanol for fungal staining. Belowground biomass was acquired using a series of sieves and water to wash away associated soil from the roots and rhizomes (Roots Lab^1^). Belowground biomass was then oven dried at 65°C to constant mass to obtain dry biomass measurements.

### Fungal assessment

Roots were stained for microscopic detection of endophytic fungi, specifically arbuscular mycorrhizal fungi (AMF) and dark septate endophytes (DSE), using the protocol described by Penn State’s Roots Lab (Roots Lab^2^). The degree of endophytic colonization was determined using a modified version of the “Slide Method” described in Giovanetti & Mosse (1980). Briefly, each sample of lateral roots was cut into five 1-cm long segments and on a 20 × 20 mm grid slide which helps in evaluating the presence or absence of AMF hyphae/vesicles/arbuscules as well as DSE hyphae/microsclerotia within each 1 mm x 1 mm square. All results are reported as % of observations with DSE presence since no AMF structures were observed.

### Data analyses

To assess the effect of whether soil conditioning species influenced the growth of our plant species (native/ introduced *P. australis*, and *S. patens*) we employed one-way ANOVA within each species, using total biomass (above plus belowground biomass) as the main response variable. Non-parametric analyses (Kruskal-Wallis one-way analysis of variance) were used if assumptions of normality (Shapiro-Wilk test) and homogeneity of variances (Bartlett’s test) were not met. One-way ANOVA’s were also used to test a variety of other response variables such as shoot emergence rate and shoot elongation rate. Two-way ANOVA was used to assess the effect of species (native/ introduced *P. australis*, and *S. patens*), soil conditioning species, and their interaction on the various response variables. Fungal colonization and BGB:AGB ratios could not be assessed using two-way ANOVA because the data was not normally distributed, so separate one-way ANOVA’s were performed to assess effect of plant species and soil conditioning species on these response variables. If our plant species experienced significant variations in biomass between soil conditioning treatments, as confirmed by the one-way ANOVA test/ Kruskal-Wallis test, then a Tukey’s HSD test was performed. Tukey’s HSD test was used to elucidate which soil conditioning treatments were significantly different from other treatments and by comparing mean values within each treatment the direction and magnitude of the soil conditioning can be assessed to confirm or reject our hypotheses. Aside from total biomass, Tukey’s HSD tests were used to assess differences between plant species and soil conditioning treatment in other analyses and traits (SIR, fungal colonization, shoot elongation/ emergence, and BGB:AGB). Tukey’s HSD tests were performed using the function “HSD.test” in R package “agricolae” (Mendiburu 2010). All figures were created using the R package “ggplot2” (Wickham 2009). While the main response variable in this study is total biomass, which we deem to be a proxy for overall plant performance, the other response variables tested provide insight into possible mechanisms of action that ultimately affect variations in biomass in plants subject to various soil communities. All data was analyzed in R studio (Version 1.1.414).

## RESULTS

### Total biomass

Soil conditioning treatment affected total biomass for both native *P. australis* (Chi-squared = 12.12, p-value = <0.01 df = 3) and *S. patens* (F_3,75_ = 2.75, p-value = 0.05) but not for introduced *P. australis* (F_3,76_ = 1.86, p-value = 0.14). Introduced *P. australis* grown in the *S. patens-*conditioned soil grew 31% smaller (19.20 g ± 8.14) than plants grown in self-conditioned soils (27.51 g ± 6.75) soils and 38% smaller than plants grown in native *P. australis-*conditioned soils (30.57 g ± 8.36) (Fig. 3, Table 3). *S. patens* grown in sterilized soils (12.65 g ± 7.32) grew 50% smaller than *S. patens* grown in introduced *P. australis-*conditioned soils, although growth differences between biotic soils differed less (self-conditioned: 18.88 g ± 5.06, native *P. australis*: 24.55 g ± 9.34, introduced *P. australis*: 25.31 g ± 7.32) (Fig. 3, Table 3). Interestingly, native *P. australis* grew the largest in introduced *P. australis-*conditioned soils (18.35 g ± 2.00) and grew 21% and 25% smaller in self-conditioned (14.49 g ± 2.64) and *S. patens-*conditioned soils (13.83 g ± 3.76) respectively (Fig. 3, Table 3).

### Individual plant-soil feedback

Introduced *P. australis* displayed positive plant-soil feedbacks when grown in soil conditioned by *S. patens*, meaning biomass was higher when grown in “home” soil than in *S. patens* conditioned soil (PSF_sp_ = 0.36 ± 0.07) (Fig. 2). Introduced *P. australis* grew slightly better in native *P. australis* than “home” soil (PSF_paa_ = -0.11 ± 0.05) as well as in sterilized soil (PSF_ster_ = -0.02 ± 0.05) (Fig. 2), suggesting that its “home” soil community was not overly pathogenic. Interestingly, native *P. australis* grew exceedingly well in introduced *P. australis-*conditioned soil (PSF_pa_ = -0.24 ± 0.03) and in sterilized soil (PSF_ster_ = -0.26 ± 0.04) than in its “home” soil; and like introduced *P. australis*, grew poorly in soil conditioned by *S. patens* (PSF_sp_ = 0.05 ± 0.05) (Fig. 2). This suggests that native *P. australis* soil communities may be dominated by pathogens. Additionally, the same microbes that illicit negative growth effects in introduced *P. australis* seem to also affect the native lineage. *S. patens* experienced much greater growth in both *P. australis-*conditioned soils (PSF_paa_ = -0.26 ± 0.06, PSF_pa_ = -0.29 ± 0.05) than in its “home soil” (Fig. 2). However, growth in sterilized soil was much lower than in its “home soil” (PSF_ster_ = 0.40 ± 0.12) (Fig. 2), suggesting that it benefits greatly from soil microbial mutualisms, especially in both *P. australis* soils.

**Fig.2.**
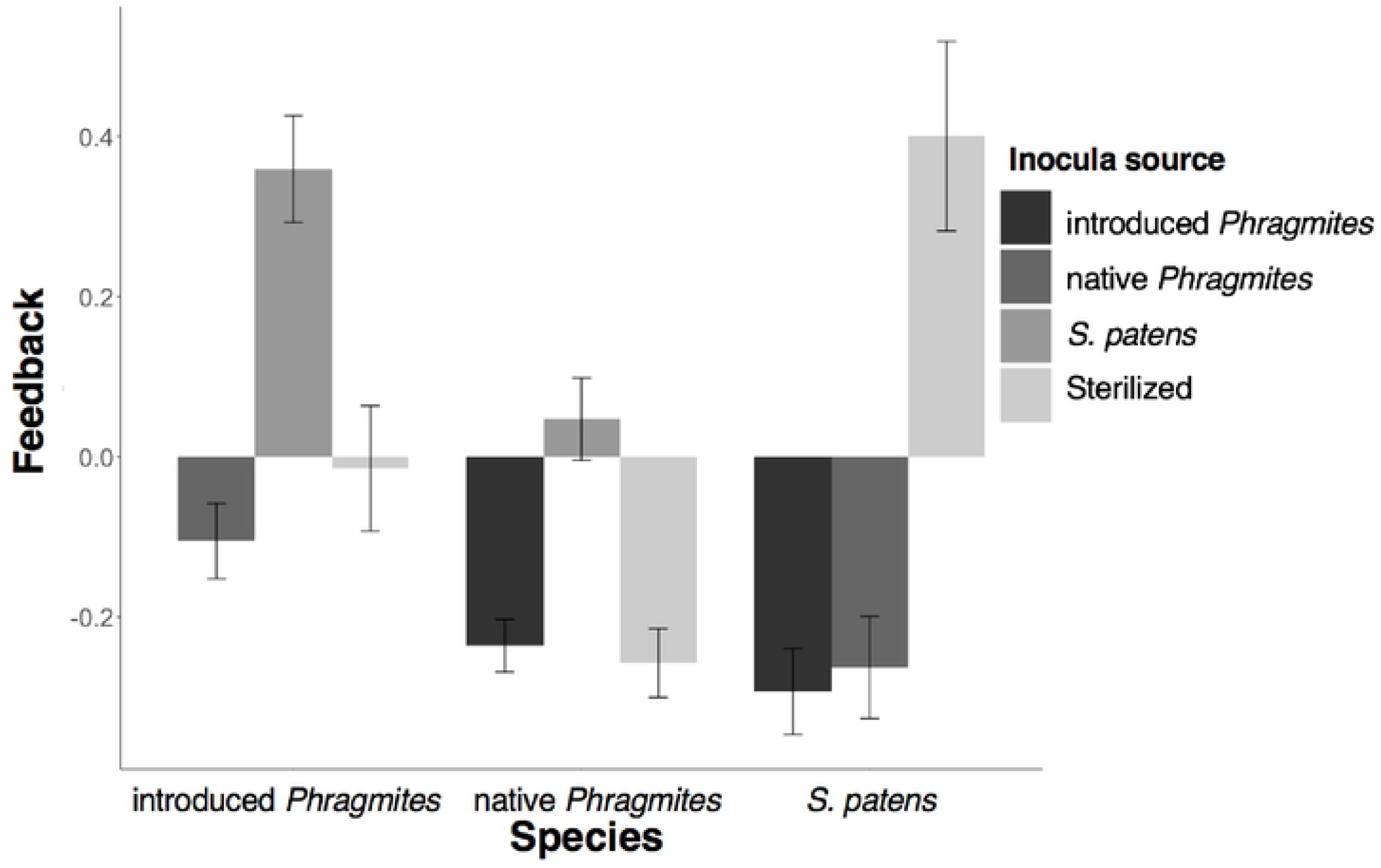
Feedbacks of introduced *P. australis*, native *P. australis*. and *S. patens* comparing pcrfonnance in conspccific soils versus hetcrospccificand sterilized soils. Positive feedback values represent superior growth in conspccific soils and negative feedback values represent superior growth in the heterospccificsoil. Error bars represent propagated ± SE. ^a^ Darkgrey = feedback of species in introduced *P. austratis* inocula, second darkestgrey = feedback of species in native *P. austratis* inocula, second light.est grey = feedback of species in *S. patens inocula*, light grey = feedback of species in sterilized soil.

**Fig.3.**
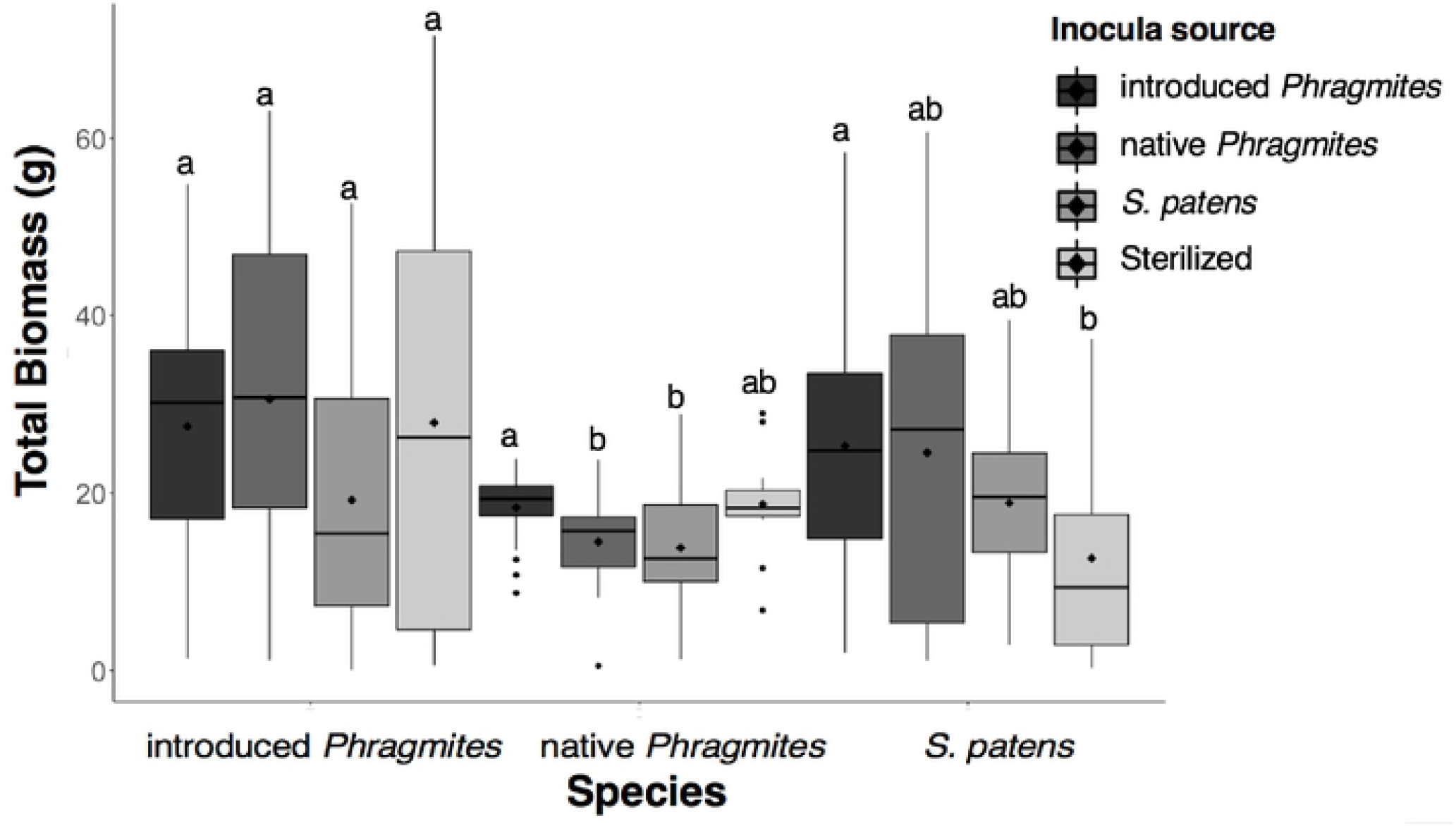
Boxplots of total biomass of introduced *P. australis*, native *P. australis*, and S. *patens* in various soil inocula sources. Black point within each boxplot represents the mean value. Upper and lower edges of each box represent the 25^th^ and 75^th^ percentile. Middle line represents the median value and whiskers represent 1.5*(25^th^ and 75^th^ quantile). Outliers arc represented by black points. ^a^Dark grey= biomass ofspecies in introduced *P. australis* inocula, second darkestgrey= biomass ofspecies in native *P. australis* inocula, second lightest grey =biomassofspecies in *S. patens* inocula, light grey = biomass of species in sterilized soil.

### Substrate-induced respiration (SIR)

The rates of soil respiration differed significantly between plant species (Table 2). Soil-conditioning species was not a strong determinant of soil respiration across species (Table 2). The interaction of species x soil-conditioning species was marginally significant in explaining soil respiration rates (Table 2). Soils associated with introduced *P. australis* (257.82 μg C-CO_2_ g soil^−1^ h^−1^ ± 132.25) respired 59% less than soils associated with native *P. australis* (625.76 μg C-CO_2_ g soil^−1^ h^−1^ ± 164.73) and 56% less than soils associated with *S. patens* (581.22 μg C-CO_2_ g soil^−1^ h^−1^ ± 298.01). Interestingly, the soils associated with introduced *P. australis* that were inoculated with conditioned soil (introduced/native *P. australis & S. patens*) respired much less than sterilized soil (Fig. 4). This suggests lower microbial biomass or community suppression in introduced *P. australis* associated soils, especially in the pots inoculated with conditioned soil. Lastly, the variation in soil respiration rates was relatively high in *S. patens* associated soils compared to the other two species (Fig. 4).

**Table 2.** ANOVA p-values of various tested response variables. The use of parametric or non-parametric tests on each factor is indicated by the color (np= non-parametric, p= parametric). Significant p-values (p-value < 0.05) indicated in bold.

|  | introduced<br>*Phragmites australis*<br>x<br>Soil Source | native<br>*Phragmites australis*<br>x<br>Soil Source | *Spartina*<br>*patens*<br>x<br>Soil Source |
| --- | --- | --- | --- |
| Total biomass | 0.14<sup>p</sup> | **<0.01<sup>np</sup>** | **0.05<sup>p</sup>** |
| Shoot elongation rate | **<0.01<sup>p</sup>** | 0.12<sup>p</sup> | 0.13<sup>p</sup> |
| Shoot emergence rate | 0.58<sup>p</sup> | 0.15<sup>p</sup> | 0.38<sup>p</sup> |
|  | Soil Source | Species | Species<br>x<br>Soil Source |
| Substrate-induced respiration<br>(SIR) | 0.29<sup>p</sup> | **<0.001<sup>p</sup>** | **0.03<sup>p</sup>** |
| BGB: AGB | 0.10<sup>np</sup> | **<0.001<sup>np</sup>** | N/A |
| Fungal colonization | 0.39<sup>np</sup> | **0.03<sup>np</sup>** | N/A |

<sup>a</sup> BGB: AGB denotes the ratio of belowground biomass to aboveground biomass. <sup>b</sup> Parametric ANOVA CRD was performed on single factors that had passed assumption testing indicated by the superscript (p). <sup>c</sup> Non-parametric Kruskal-Wallis test was performed on single factors that failed assumption testing indicated by the superscript (np). <sup>d</sup> Factorial analyses were performed using two-way ANOVA. N/A denotes factor and variable combinations not tested.

**Table 3.** Response variable means and standard error (SE±) grouped by soil inocula source within each species. Tukey’s HSO test was performed within each species.

| Species | Soil | Biomass |  |  | SIR |  |  | BGB: AGB |  |  |
| --- | --- | --- | --- | --- | --- | --- | --- | --- | --- | --- |
| | | Mean (g) | SE (±) | HSD | Mean | SE (±) | HSD | Mean | SE (±) | HSD |
| introduced<br>*Phragmites australis* | PA | 27.51 | 6.75 | a | 227.15 | 63.00 | cd | 0.65 | 0.12 | a |
|  | PAA | 30.57 | 8.36 | a | 262.10 | 35.05 | bcd | 0.72 | 0.18 | a |
|  | SP | 19.20 | 8.14 | a | 181.64 | 43.76 | d | 0.64 | 0.23 | a |
|  | STER | 27.92 | 11.96 | a | 463.01 | 64.94 | abcd | 0.53 | 0.16 | a |
| native<br>*Phragmites australis* | PA | 18.35 | 2.00 | a | 708.50 | 94.54 | a | 0.44 | 0.06 | a |
|  | PAA | 14.49 | 2.64 | b | 541.20 | 44.46 | abc | 0.43 | 0.06 | a |
|  | SP | 13.83 | 3.76 | b | 606.95 | 81.86 | ab | 0.39 | 0.07 | a |
|  | STER | 18.75 | 3.02 | ab | 666.99 | 99.11 | ab | 0.41 | 0.05 | a |
| *Spartina patens* | PA | 25.31 | 7.32 | a | 667.37 | 97.93 | a | 0.29 | 0.09 | a |
|  | PAA | 24.55 | 9.34 | ab | 635.37 | 163.89 | ab | 0.18 | 0.07 | ab |
|  | SP | 18.88 | 5.06 | ab | 541.41 | 196.82 | abc | 0.24 | 0.06 | ab |
|  | STER | 12.65 | 5.70 | b | 380.26 | 54.46 | abcd | 0.15 | 0.02 | b |
| Species | Soil | % Fungal colonization |  |  | Shoot Elongation rate |  |  | Shoot emergence rate |  |  |
| | | Mean | SE (±) | HSD | Mean | SE (±) | HSD | Mean | SE (±) | HSD |
| introduced<br>*Phragmites australis* | PA | 21.50 | 26.35 | a | 7.81 | 0.69 | a | 0.08 | 0.02 | a |
|  | PAA | 26.00 | 35.85 | a | 6.31 | 0.68 | ab | 0.07 | 0.04 | a |
|  | SP | 2.50 | 2.56 | a | 6.06 | 0.98 | ab | 0.07 | 0.03 | a |
|  | STER | 3.50 | 7.00 | a | 3.48 | 0.74 | b | 0.05 | 0.04 | a |
| native<br>*Phragmites australis* | PA | 6.00 | 15.01 | a | 4.03 | 0.65 | a | -0.02 | 0.01 | a |
|  | PAA | 3.00 | 5.66 | a | 3.63 | 0.73 | a | -0.02 | 0.01 | a |
|  | SP | 0.22 | 0.67 | a | 3.06 | 1.29 | a | -0.01 | 0.01 | a |
|  | STER | 2.00 | 4.00 | a | 2.56 | 1.01 | a | -0.02 | 0.01 | a |
| *Spartina patens* | PA | 13.25 | 31.07 | a | 4.89 | 1.62 | a | 0.17 | 0.08 | a |
|  | PAA | 1.25 | 2.38 | a | 5.22 | 1.85 | a | 0.19 | 0.09 | a |
|  | SP | 4.00 | 5.76 | a | 6.72 | 1.34 | a | 0.18 | 0.07 | a |
|  | STER | 3.00 | 6.00 | a | 4.58 | 1.30 | a | 0.10 | 0.07 | a |

<sup>a</sup> Units: Biomass= grams (g), Substrate-induced respiration (SIR)= $\mu\text{g C-CO}_2 \text{ g soil}^{-1} \text{ h}^{-1}$ , Shoot elongation= mm per day<sup>-1</sup>, Shoot emergence= # of shoots per day<sup>-1</sup>

**Fig.4.**
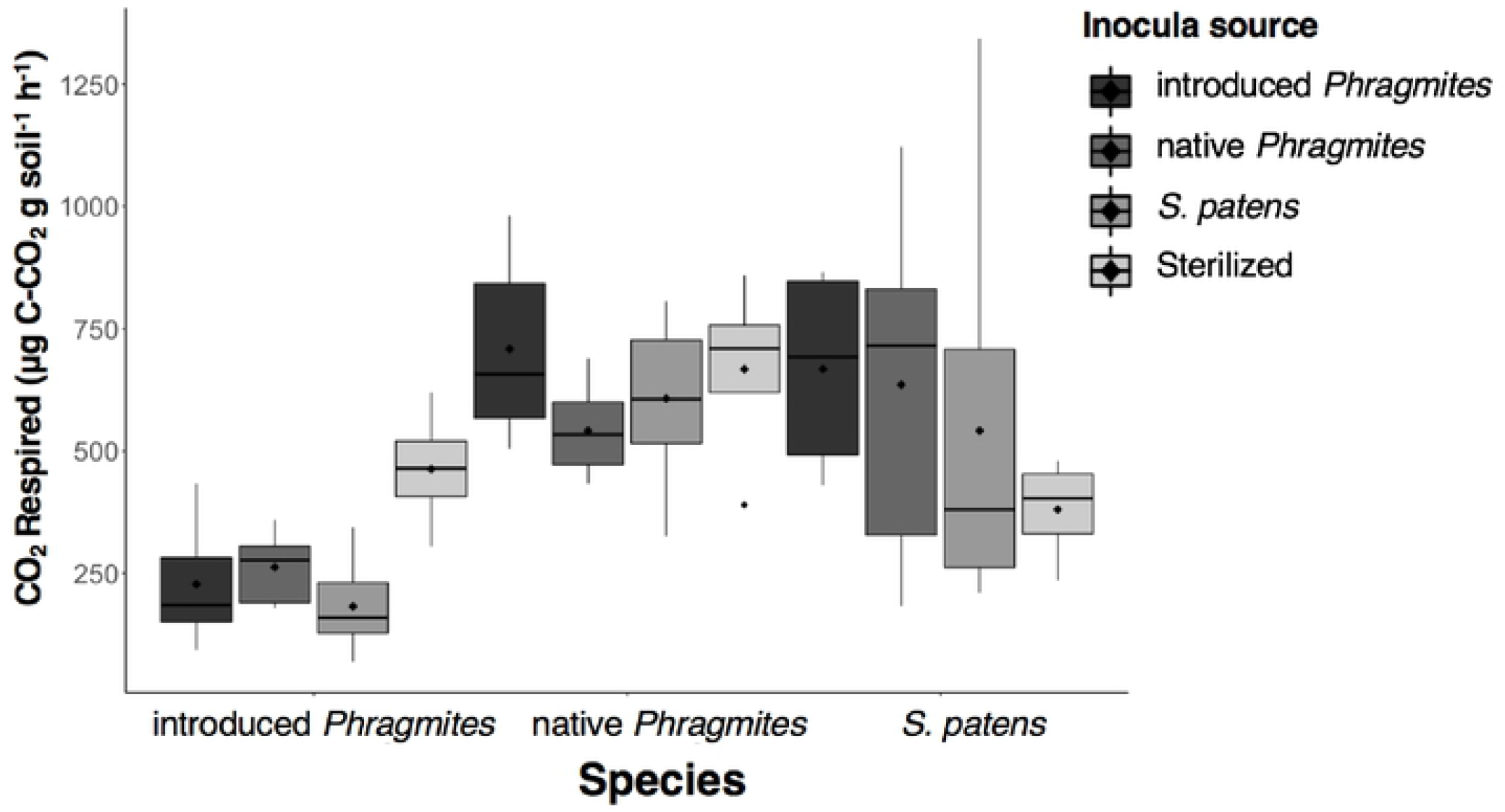
Boxplots of substrate-induced respiration of introduced *P. australis*, native *P. australis*, and S. *patens* in various soil inocula sources. Black point within each boxplol represents the mean value. Upper and lower hinges of each boxplot represent the 25^th^and 75^th^ pereentile. Middle line represents the median value and whiskers represent 1.5* (25^th^ and 75^th^ quantile). Outliers are represented by black points. ^a^Dark grey = SIR of species in introduced *P. australis* inocula, second dark est grey = SIR of species in native *P. australis* inocula, second lightest grey = SIR of species in *S. patens* inocula, light grey = SIR of species in sterilized soil.

### Fungal assessment

All root endophytes identified belonged to the group DSE and no structures resembling AMF were found in any of our samples. The variation in root colonization rates was most evident between plant species, with introduced *P. australis* roots showing the highest rate of colonization with 15.17% ± 25.73 of the root length colonized compared to only 2.71% ± 8.08 in native *P. australis* and 5.71% ± 17.00 in *S. patens* (Fig. S1). Soil conditioning treatment on the other hand, was not significant in explaining fungal colonization frequency (Table 2). Although, there does seem to be a higher rate of colonization of introduced *P. australis* roots specifically in native (26.00% ± 35.84) and introduced (21.50% ± 26.45) *P. australis-*conditioned soils (Fig. S1).

### Shoot emergence and elongation rate

Soil conditioning species had a large effect on the shoot elongation rate of introduced *P. australis* (Table 2), with plants grown in “home” soil (7.81 mm per day^-1^ ± 0.69) growing 56% faster than plants grown in sterilized soil (3.48 mm per day^-1^ ± 0.74) and marginally faster than native *P. australis* soil (6.31 mm per day^-1^ ± 0.68) at 19% and *S. patens* soil (6.06 mm per day^-1^ ± 0.98) at 22% (Fig. S3, Table 3). Variation in shoot elongation rate in native *P. australis* and *S. patens* did not differ between soil-conditioning species (Table 2). The rate of shoot emergence was not strongly influenced by soil-conditioning species in any of our species (Table 2). Admittedly, many of the plants experienced pot binding and lodging by the end of the growth period which could be confounding variables obscuring shoot emergence rate. There does seem to be a trend of *S. patens* and introduced *P. australis* having lower shoot emergence rates in sterilized soil (Table 2); with introduced *P. australis* grown in sterilized soil (0.05 shoots per day^-1^ ± 0.04) having 37.5% lower emergence rates than plants grown in “home” soil (0.08 shoots per day^- 1^ ± 0.02) (Fig. S2, Table 3). This trend also holds true for *S. patens* as plants grown in sterilized soil (0.10 shoots per day^-1^ ± 0.07) have 45% lower emergence rates than plants grown in “home” soil (0.18 shoots per day^-1^ ± 0.07) (Fig. S2, Table 3).

### Seedling mortality

Seedling mortality was only observed in *S. patens*, with a total of 71 *S. patens* seedling deaths during the growth period (Table S1); often multiple sequential deaths and replacements occurred within a single replicate. A Poisson regression revealed that seedling mortality was not significantly affected by soil treatment (z-value = 1.73, Pr(>|z|) = 0.08), although *S. patens* grown in “home” soil tended to experience 50% greater seedling mortality (26 deaths) than plants grown in sterilized soil (13 deaths) (Table S1). There also seemed to be lower incidence of seedling death when *S. patens* was grown in both *P. australis-*conditioned soils as plants grown in “home” soil experienced 47% and 39% greater seedling mortality than plants grown in native and introduced *P. australis* respectively (Table S1). No seedling death was observed in native or introduced *P. australis* seedlings.

## DISCUSSION

In this study, we characterized the PSF profiles of introduced and native *P. australis* as well as the common saltmarsh grass, *S. patens*. As we hypothesized (H1), introduced *P. australis* exhibited a neutral conspecific PSF indicating that rhizosphere soil communities neither directly hinder nor promote the invasive spread of *P. australis*. Our results corroborate Bowen et al. (2017), which found no direct influence of microbial communities on the biomass of introduced *P. australis*. Given the simultaneous spread of introduced *P. australis* and the decline of native *P. australis* and *S. patens*, we hypothesized (H2) that native plants would have a negative conspecific PSF (negative growth effects in self-conditioned soil communities). Our findings supported H2 as negative conspecific feedbacks were observed in native *P. australis*. Allen et al. (2018) found negative conspecific feedbacks in both native and introduced *P. australi*s while our results only indicate negative conspecific feedbacks in the native lineage. Unlike previous studies, we found that introduced *P. australis* can have strong positive impacts on the growth of *S. patens* through positive heterospecific PSFs. Our research indicates that PSF interactions between these two dominant species in wetlands may differ from what has been predicted in the literature (Amsberry et al. 2000; Minchinton & Bertness 2003; Lonard et al. 2010).

Contrary to our hypothesis (H1), *S. patens* experienced positive-growth effects when grown in soil inoculated with introduced *P. australis-*conditioned soil inoculum (Fig. 2). In the reciprocal treatment, the growth of introduced *P. australis* declined when grown in *S. patens-*conditioned soil incoclum, suggesting an antagonistic effect of *S. patens*-associated soil community to introduced *P. australis* growth (Fig. 2). Previous studies have shown that *S. patens* can impede *P. australis* invasion in the field, which may in part result from *S. patens* soil pathogen community spillover effects on *P. australis* growth as indicated by the negative effect of *S. patens* inocula on *P. australis* growth (Wang et al. 2006; Kettenring et al. 2015). Accounts of disturbance-mediated spread of introduced *P. australis* into *S. patens-*dominated wetlands shaped our initial supposition that introduced *P. australis-*conditioned soil would have an antagonistic effect on *S. patens* (Amsberry et al. 2000; Minchinton & Bertness 2003; Lonard et al. 2010). However, our data suggest these disturbance events not only aid in reducing local plant competition for introduced *P. australis* but reduce the load of *S. patens* associated soil microbes allowing for introduced *P. australis* seedling recruitment (Wang et al. 2006; Kettenring et al. 2015).

### Intraspecific variability in plant-soil feedbacks of *P. australis*

Our data support H1, neutral conspecific PSFs in introduced *P. australis* (growth in conspecific ∼ sterilized), and H2, negative conspecific PSF in native *P. australis* (growth in conspecific < sterilized, Figs. 2-3). Thus, plant-soil interactions negatively affected native *P. australis* but not introduced *P. australis*. These findings agree with the few other intraspecific PSF studies, which have found intraspecific variation in the PSFs of the model species *Arabidopsis thaliana*, leguminous perennial *Trifolium pratense*, as well as the North American invasive *Ailanthus altissima* (Felker-Quinn et al. 2011; Bukowski & Petermann 2014; Wagg et al. 2015; Bukowski et al. 2018). Another PSF study of *P. australis* found no difference in feedbacks between native and introduced lineages as both were negative (Allen et al. 2018). Several factors may have caused the differences in results between ours and Allen et al. (2018) including differences in salinities, nutrient level, etc. between the two studies as well as local variability in soil microbial communities used in the initial inocula. Unlike the native and Gulf lineage of *P. australis*, introduced *P. australis* can exhibit high site-to-site variability in soil microbial community composition (Bowen et al. 2017). Differences in soil communities between *P. australis* lineages may underlie some of the differences in plant performance and invasive success of some lineages over others.

Our design allowed for comparison of PSFs of native and introduced *P. australis* in each other’s soil inoculum. While the introduced *P. australis* experienced no significant variation in biomass between native versus introduced soil inocula, the native *P. australis* produced greater biomass when grown in introduced *P. australis* inocula (Figs 2-3). The increased biomass of native *P. australis* in introduced *P. australis* inocula may be due to accumulation of species-specific soil pathogens within “home” soil communities. Growth of native *P. australis* in heterospecific soils may therefore be greater due to lower antagonism from specialized pathogens. This finding supports the Janzen-Connell hypothesis which posits that progeny tend to establish farther from parents due to lower density of specialist pathogens/herbivores (Janzen 1970; Connell 1971). Bowen et al. (2017) reveals high conservation in soil community in native *P. australis* across different populations, which, taken with our results, suggests that a relatively high portion of these conserved microbes may be pathogens. Conversely, there may not be more harmful pathogens in native *P. australis* soil communities but there may be protective/mutualistic microbes found in the soil communities of introduced *P. australis*. Some fungal endophytes found in introduced *P. australis* can protect seedlings from pathogens and promote survival and growth as well as increase ability to deal with abiotic stressors (Kowalski et al. 2015; Shearin et al. 2018; White et al. 2018; Gonzalez Mateu et al. 2020). Additionally, there is some evidence that introduced *P. australis* suffers less from foliar pathogens and this may also be the case with soil pathogens, which explains why there was less variation in growth between both *P. australis* inocula observed in introduced *P. australis* compared to the native lineage (Allen et al. 2020).

Our results demonstrate variations in intraspecific PSFs within and between lineages which may inform the difference in success between native and introduced *P. australis*. Release from enemies often results in enhanced growth as predicted by the Enemy-Release Hypothesis, which was originally used to explain the success of introduced species and holds for soil inocula effects from varying species (Keane & Crawley 2002). Other studies have found that, while native grasses tend to cultivate negative intraspecific feedbacks, invasive grasses tend to have neutral or positive intraspecific PSF (Kulmatiski et al. 2008; Hawkes et al. 2013). Accordingly, native *P. australis* like other native grasses tended to cultivate a negative feedback in “home” soil (Fig. 2). Comparisons of native versus introduced *P. australis* have identified highly pathogenic strains of *Pythium spp*. found in native *P. australis* dominated sites which may explain the negative PSFs seen in the natives (Crocker et al. 2015; Crocker et al. 2017). Interestingly, we found that fungal colonization by DSE in introduced *P. australis* inoculated with native and introduced conditioned soil was unusually high (Fig. S1). This result in line with Gonzalez Mateu et al. (2020), which found that introduced *P. australis* inoculated with DSE strains performed better in mesohaline (0.5-5 ppt) conditions, suggesting that high colonization rates by DSE may partially explain we observed neutral PSFs in introduced *P. australis*. While differences in physiology and growth pattern may explain the success of introduced *P. australis* relative to native *P. australis*, our results provide the first discrete evidence that negative PSFs may play a role in reducing native *P. australis* competitiveness against introduced *P. australis* invasion.

### Interspecific soil conditioning effects of *P. australis*

Contrary to our initial hypothesis (H1), *S. patens* experienced no negative growth effects from introduced *P. australis* inocula. Surprisingly, *S. patens* growth tended to increase with introduced *P. australis* inocula (Figs 2-3). Seedling death of *S. patens* was still observed in both introduced and native *P. australis*-conditioned soils, however *S. patens* seedling death was observed in every soil treatment and tended to be higher in *S. patens*-conditioned soils (Table S1). Variation in seedling mortality between different species is well documented in upland plants and stems from tradeoffs between survivability of individual propagules and propagule output per plant (Canham et al. 1999; Moles & Westoby 2004; Milbau et al. 2017). Therefore, the variation in mortality between the different species in this study likely stems from propagule variation rather than soil conditioning treatment. These results counter past literature that has generally found negative plant growth effects associated with introduced *P. australis* soil communities. Multiple studies have shown introduced *P. australis* elicits negative growth responses in native plants through soil community assemblages or allelopathy, while native *P. australis* has been shown to promote native species growth (Crocker et al. 2017; Uddin et al. 2017; Allen et al. 2018). Many invasive plants cultivate soil communities that have variable growth effects in native plant communities; while some plants may experience negative growth effects when grown in invasive-conditioned soils, other native species may experience positive growth effects (Yelenik & Levine 2011; Shannon et al. 2012; Perkins & Nowak 2013). Our data represent the first evidence of growth facilitation of introduced *P. australis* inocula on the growth of native plants.

Our SIR analysis revealed that introduced *P. australis* had lower active microbial biomass in surface soils (Fig. 4). Our results support the Burke et al. (2002) findings in that our introduced *P. australis* soils had a nearly two-fold reduction in active microbial biomass compared to *S. patens* soils. Bernal et al. (2017) similarly demonstrated that *P. australis* surface soils tended to respire less than deeper soils, an unusual pattern for most soils, which normally exhibit the greatest microbial biomass and activity at the surface (Fang & Moncrieff 2005). Although surface soil respiration is lower in introduced *P. australis* compared to the native lineage or *S. patens*, deep soil priming effects may offset low surface soil respiration (Bernal et al. 2017). The deep soil priming effects of introduced *P. australis* suggest that it may be exuding more labile C in deeper soil depths than in shallower soils, which has been shown to have lower surface soil signatures of bacterial PLFAs than other species such as *S. alterniflora* (Zhang et al. 2017^2^). Introduced *P. australis* can send roots to depths exceeding 3 m, whereas *S. patens* rooting-depth is confined to the top 30 cm of soil (Windham 2001; Meschter 2015). The fact that introduced *P. australis* and *S. patens* inhabit different depths along the soil column in turn influencing the amount of microbial biomass in these different strata may inform the interspecific PSFs observed in this study as we only test PSFs in surface soil, and as a result ignore deep soil microbial communities.

Taken together, our results contribute to the growing body of evidence that *S. patens* exhibits biotic resistance against introduced *P. australis* invasion (Figs 2-3). Several studies demonstrate the ability of *S. patens* to compete against introduced *P. australis*, often existing in competitive standoff when the integrity of meadows is maintained, and invasion *S. patens* by introduced *P. australis* is only seen in instances of disturbance (Burdick & Konisky 2003; Wang et al. 2006; Kettenring et al. 2015). Despite the negative growth effects of *S. patens* soil inoculum on *P. australis*, expansion of *P. australis* is often observed to occur through vegetative growth of rhizomes and stolons which may be less susceptible to negative PSFs than seedlings (Burdick et al. 2001). Even negative interspecific PSFs may favor *S. patens*, the effects of shading imposed by the tall shoots of introduced *P. australis* along the invasion front may shift competitive interactions against *S. patens*. It is worth noting that introduced *P. australis* commonly establishes in areas of disturbance, where *S. patens* is not present, and perhaps soil legacy effects degrade rapidly following disturbance, which may allow *P. australis* seedling to establish (Minchinton & Bertness 2003; Kulmatiski & Kardol 2008; Kettenring et al. 2015). Following this disturbance, vigorous vegetative *P. australis* can advance into the disturbed area, and seeds may be able to establish in the newly vacant soil, thus overwhelming any advantage *S. patens* has belowground (Meyerson et al. 2000; Minchinton & Bertness 2003; Kettenring & Whigham 2018). *S. patens* has innate physiological and morphological characteristics that aid in competition against invasive plants, and our results demonstrate that soil microbial community may contribute to this observed biotic resistance against *P. australis* invasion into native wetlands. However, maintaining the integrity of native wetlands and limiting instances of disturbance is key in curbing the invasion of introduced *P. australis*.

### Limitations and future direction

Most plant-soil feedback studies have been performed in upland habitats, but our understanding of coastal wetland PSF is growing, and results have been conflicting. This study advances our knowledge of the specific variables that dictate PSFs in wetland systems (DSE’s, microbial biomass, native vs. non-native species), as they may differ considerably from uplands. Salinity, water level flux, soil aeration, and nutrient fluctuations are all variables that exert strong control over soil community and physiological tolerances of the plant. Any of these factors could modify PSFs in wetlands, and we suggest that future coastal wetland PSF studies investigate natural gradients in these factors. Furthermore, future studies should include higher replication in soil and seed populations than was possible herein. The effect of anthropogenic forces important in the spread of introduced *P. australis*, such as nutrient pollution and physical disturbance, should be evaluated in the context of PSF to examine whether these forces color the context of PSFs and in turn influence introduced *P. australis* spread.

## ACKNOWLEDGEMENTS

We gratefully acknowledge the help given to us in sourcing and collecting our field samples by the Smithsonian Environmental Research Center (SERC; Edgewater, MD). Native *P. australis* seed was graciously sourced and collected by help from Kurt Kowalski, Ph.D. (United States Geological Survey, USGS). USFWS Research & Monitoring Special Use Permit #2018014 was used for Cedar Point National Wildlife Refuge sampling. Funding assistance was provided by Villanova University and sampling at SERC was supported by the National Science Foundation Long-Term Research in Environmental Biology Program (DEB-0950080, DEB-1457100, and DEB-1557009).

## AVAILABILITY OF DATA AND MATERIALS

All data used to run the data analyses are available at the following link (https://github.com/seanlee218/Phragmites_PSF_Study_data.git).

## AUTHOR CONTRIBUTIONS

S.L., J.A.L., and S.C. in conjunction designed the experiment. A.H.B. and M.G.M. helped with sampling of *P. australis* material. All procedures and data collection were performed by S.L. Guidance and assistance in seed germination was offered by T.M. Data analyses were performed by S.L. First draft was written by S.L. and all authors edited the manuscript.

